# Structural study of membrane proteins using vesicles

**DOI:** 10.1101/2023.06.17.545446

**Authors:** Hang Liu, Shangyu Dang

## Abstract

Membrane proteins play crucial roles in numerous biological processes and are important drug targets. However, structural studies of memebrane proteins heavily rely on solubilization by detergents, which may not reflect their native states in the cellular context. Moreover, identifying suitable detergents for individual membrane proteins is a tedious and costly screening process. Here, we developed a vesicle-based method that enables membrane protein structure determination in their native lipid environment, thereby bypassing the limitations of detergent solubilization. Using this approach, we isolated vesicles containing the multidrug efflux transporter AcrB and solved its structure by cryo-electron microscopy. Intriguingly, the AcrB trimer in the vesicle exhibited a loose assembly compared to the detergent-solubilized and nanoparticle structures. Our method presents a promising approach for studying structure and function of membrane protein in their native environment without the need for detergent screening and protein purification.

## Introduction

High-resolution structures provide valuable information to understand the working mechanism of biological macromolecules. Particularly, membrane proteins, which contribute to more than 60% drug targets^1,2^, play versatile physiological functions. Since the first atomic-resolution structure of membrane protein was determined in 1985^3^, numerous structures of membrane proteins have been captured^4^. These structures not only deepen our understanding of their molecular mechanism, but also facilitate drug development for the treatment of related diseases. Routinely, isolation of membrane proteins from the native cell membrane using detergents is required for subsequent structural and biochemical studies^5,6^. A lot of effort has been spent to obtain well-behaved membrane proteins by screening detergents since individual membrane proteins behave differently to various detergents^7–9^. Although the structures determined by using detergent-solubilized membrane proteins have advanced our understanding in the past decades, the limitations are obvious.

First, lipids, important regulators to a majority of membrane proteins^10–12^, have been omitted during detergent extraction. Thus, it is incapable of studying the lipid regulatory mechanism using the detergent system. Second, to obtain enough membrane protein suitable for biochemical assay, a lot of effort has been put into detergent screening. For membrane proteins that are sensitive to local environmental changes, purification may fail due to aggregation, unfolding or other undesirable results. Third, multiple steps, including affinity chromatography and size exclusion chromatography, have been used in membrane protein purification to ensure the high purity of the target protein for subsequent experiments. However, the weak interacting partners may be lost during thes processes, and opportunities to study the interaction of the target membrane protein and its binding proteins are missed.

In the last decade, several technologies have been developed to study membrane protein in its native environment by mimicking the lipid environment^13^. These include nanodisc^14,15^, salipro^16^, SMALPs^17^, and proteoliposome system^18–20^. However, the lipids used in these methods are usually different from the composition of those in the native cell membrane, and therefore may not represent the actual native states of these membrane proteins.

In this study, we developed a vesicle-based technology to study membrane proteins in their native membrane environment. By skipping the isolation of target membrane protein by detergent, this method maximally preserves the native lipid environment and may provide *in-vivo* information that has been lacking in previous studies. In addition, this method also saves abundant time and cost by bypassing the tedious optimization of purification, including detergent screening. The method reported here has the potential to unravel the mechanism of interested membrane proteins in their native states.

## Results

### Generation of vesicles

We chose the multidrug efflux transporter AcrB, a homotrimeric intergral membrane protein^21,22^, as an example to test the vesicle-based method. Harvested *E. coli* cells with AcrB overexpressed were treated with lysozyme to break the cell walls and release the cell membrane^23,24^. The French press was then used to facilitate the formation of inside-out vesicles. Soluble proteins and unbroken cells were removed by high-speed and ultra-speed centrifugation. The pellet was re-suspended and fractionated by sucrose gradient centrifugation to separate protein-containing vesicles from empty ones (Figure 1A). The clear band on SDS-PAGE representing AcrB indicated successful isolation of vesicles with the target protein (Figure 1B). In comparison to the sample before sucrose gradient centrifugation, the one after showed clearer background and homogeneous vesicles in size (Figure 1C-D). The majority of the vesicles were less than 100 nm or even 50 nm in size, making them suitable for cryo-EM studies (Figure 1D).

**Figure 1.**
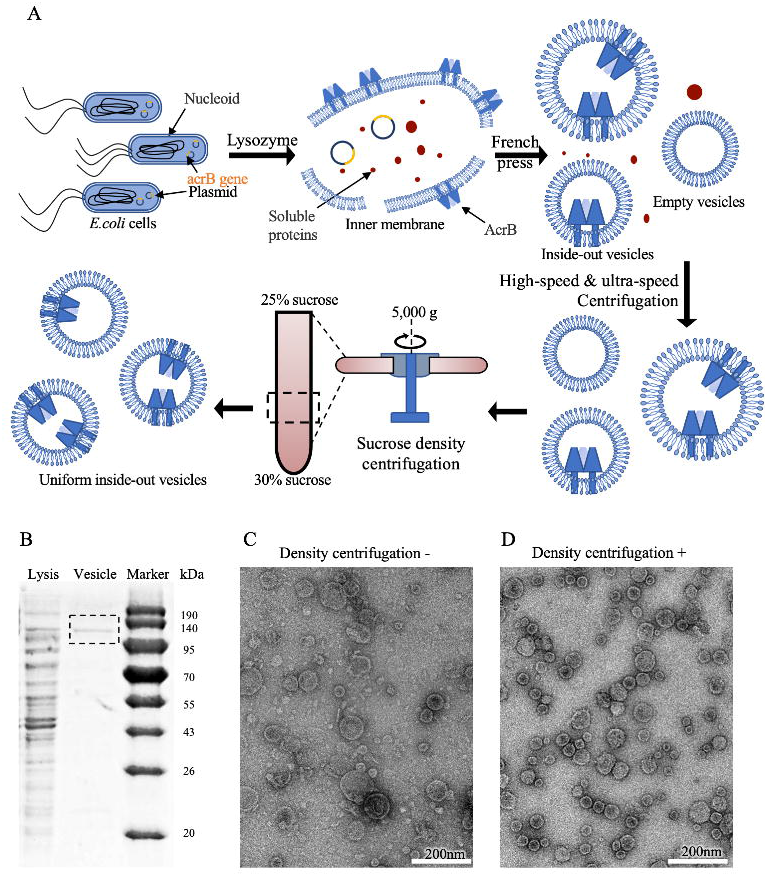
Generation of inside-out vesicles from *E. coli* cell membrane. (A)A workflow for inside-out vesicle preparation using the AcrB overexpressed bacterial cells. (B) SDS-PAGE indicates successful vesicle preparation. Lysis: cell lysate after the French press containing various vesicles and soluble proteins. Vesicle: vesicle sample after high-speed and ultra-speed centrifugation. The target band of AcrB (∼110 kDa) was indicated with a black dash box. (C) A representative negative stain image of the vesicle sample without sucrose density centrifugation purification. (D) A representative negative stain image of one uniform vesicle fraction after sucrose density centrifugation.Scale bars were indicated in the right bottom of the images.

### Structure of AcrB in vesicle

The collected vesicles were concentrated before loading onto holey carbon grids for cryo-sample preparation. The cryo-EM micrograph suggested most of the vesicles are smaller than 50 nm (Figure 2A). Particles were first picked randomly with the blob picker of cryoSPARC and sorted out by 2D classification^25^. The representative 2D averages were then used as templates to guide accurate particle picking with assistance of the template picker and Topaz^26,27^ (Figure 2B). Particles from 2D classes showing clear AcrB features were used for 3D reconstruction (Figure 2C). Based on the membrane curvature in the 3D reconstruction, the vesicle was around 450 Å in diameter by rough calculation (Figure 2C), consistent with observation from raw micrographs and 2D averages. To improve resolution of AcrB, a mask only covering the protein density was applied. In the medium resolution structure of AcrB, the domain organization of soluble regions could be observed, while transmembrane regions are too weak to be seen clearly. This medium resolution cryo-EM density map enabled us to systematically compare structures of AcrB from vesicle and nanoparticle. We individually docked three AcrB molecules into the cryo-EM density to get the corresponding AcrB homotrimer representing its native state in vesicles (Figure 2D). Intriguingly, structural comparison using one protomer as a reference indicated a loose assembly of the trimeric AcrB (Figure 2E). Specifically, the other two protomers in the vesicle structure are more distant from the reference protomer than those in the nanoparticle structure by 17 Å and 16 Å, respectively. In addition, these two protomers rotated 6 degrees against the symmetric axis of the AcrB trimer compared to structure of AcrB in nanoparticle^28^. However, limited by the resolution of AcrB in this study, it was unclear whether AcrB adopted internal conformational differences other than the trimeric assembly, which needs to be evaluated in future research. Briefly, the structure of AcrB obtained in vesicles, likely presenting its native state, reflects conformational differences to structure of AcrB obtained by either detergent solubilization or nanoparticle reconstruction. The relevance of these structural differences and physiological functions of AcrB requires further study.

**Figure 2.**
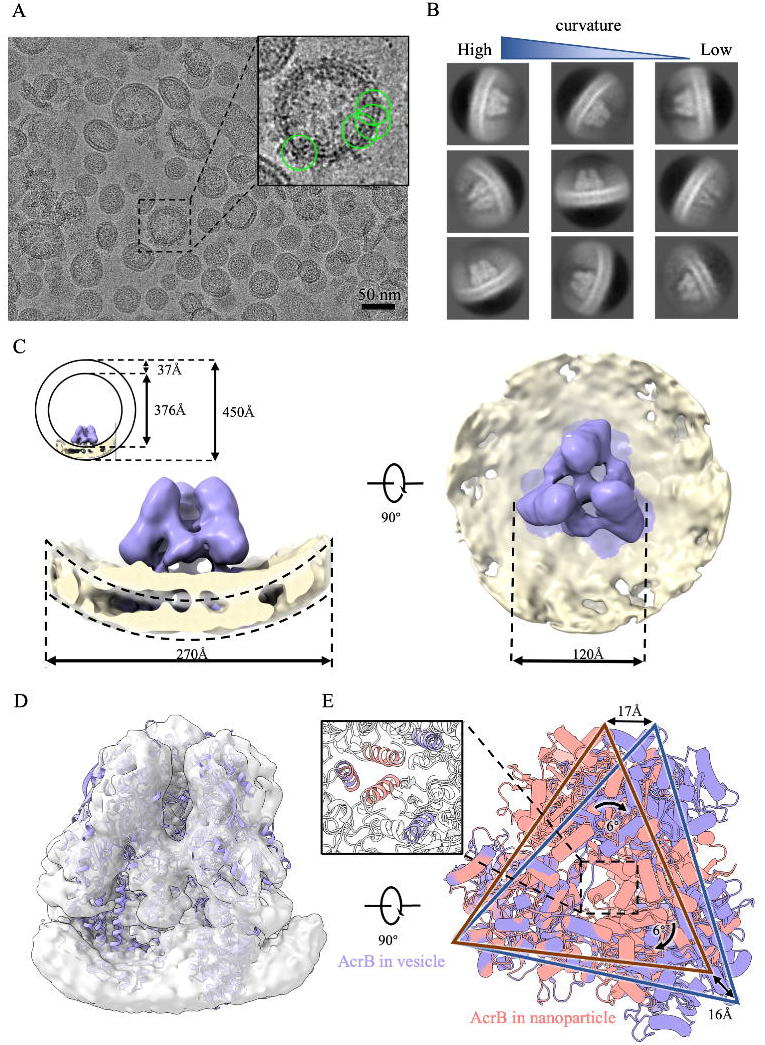
Cryo-EM structure determination of AcrB in vesicle. (A) A representative cryo-EM image of vesicles. A typical vesicle containing AcrB (green circle) was showed in the zoom-in view. (B) The representative of 2D class averages showed clear AcrB features. 2D classes showed different membrane curvatures were also indicated. (C) Side (left) and top (right) view of cryo-EM density map of AcrB (purple) in vesicle membrane (yellow). The original curvature of the vesicle membrane was shown with the black dashed lines. The average vesicle thickness and size were labeled in the upper-left. (D) Cryo-EM density map of AcrB from local refinement focusing on its soluble region. The three protomers of AcrB (purple, PDB code: 6BAJ) were independently docked into the cryo-EM map of AcrB in vesicle. (E) Structural comparison of AcrB from vesicles (purple, this study) and cell membrane nanoparticles (orange, PDB code: 6BAJ). Structural differences are highlighted with distance and rotation angle labeled.

## Discussion

In summary, here we reported a simple method to study membrane proteins using vesicles, which maximally retain their native lipid environment. Compared to previously determined structures of AcrB in detergent^29^ and nanoparticle^28^, the structure of AcrB in the vesicle exhibits a loose assembly of the trimeric complex. However, due to the resolution limitation, other detailed structural differences are yet to be identified. Additionally, the linkage between the loose assembly of AcrB in vesicle and its physiological functions requires further investigation.

The medium resolution of AcrB in this study is likely limited by the number of featured particles used for the final reconstruction. Collecting a huge dataset could help improve the resolution despite the low efficiency. Another strategy is enriching vesicles containing target protein in the cryo-grids for single-particle cryo-EM data collection, which could be achieved by adding additional steps in vesicle preparation, such as affinity chromatography, or cryo-sample preparation, such as affinity grids undermentioned. Further optimization of this method would expand its application in structural and biochemical studies of membrane proteins without detergents.

For a long time, structural studies of integral membrane proteins heavily relied on their solubilization by detergents. Undoubtedly, these structures have advanced our understanding of the working mechanism of the interested membrane proteins. To investigate the influence of the lipid environment on membrane proteins, many strategies, including nanodisc, have been developed to mimic the native lipid context. In this study, we move forward by introducing a vesicle-based method to facilitate membrane protein structure determination in their native membrane environment. With further optimization, this method could be used widely to capture the native state of interesting membrane proteins, and may provide unprecedented findings.

Compared to methods, like nanodisc, the vesicle-based method provides a relatively closed system. Additionally, the protein orientation in the vesicle membrane can be changed to be similar or opposite to its native orientation. In the vesicle preparation, the buffer condition inside or outside could be altered if needed, which would provide an opportunity to study target protein under various cellular stimuli.

In contrast to the proteoliposome system, the vesicle-based method provides a membrane environment that is more similar to the cell membrane environment in which the target membrane proteins naturally exist. The native lipid composition may stabilize protein in a physiologically relevant state, and enable us to study the regulatory mechanism of the target membrane protein by some lipids.

The size of the vesicle could be adjusted during preparation. Larger vesicle have a relatively flattened membrane surrounding target proteins, while smaller vesicles will cause the membrane surrounding target proteins to bend sharply, generating different membrane forces (Figure 3C). Previous studies have suggested that the transduction of mechanical forces through mechanical sensor proteins is mediated by membrane bending^30^. The vesicle-based method can be used to study how mechanical sensor proteins respond to forces of different magnitudes generated by membrane bending. Furthermore, water/ion channels can be co-expressed with target protein for vesicle generation, and the activation of these channels can be used to provide different osmotic pressure or ion strength prior to cryo-sample freezing to capture different states of the target membrane protein under various conditions (Figure 3C).

**Figure 3.**
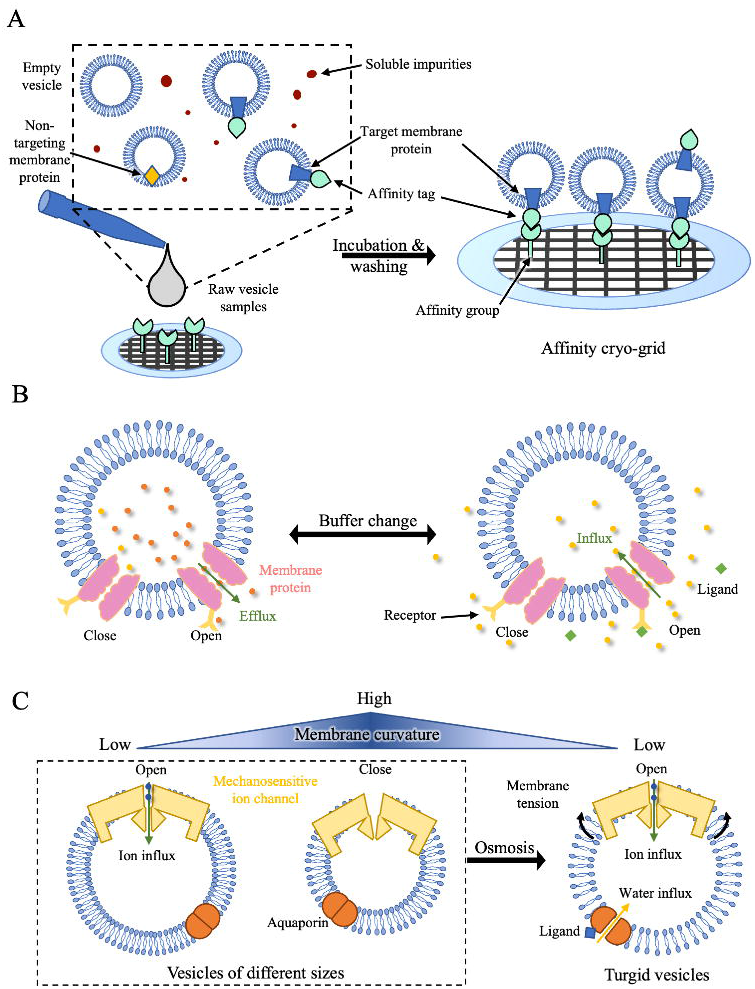
The potential application of the vesicle-based method. (A) Schematics of vesicle-based method combined with affinity cryo-grids for structural studies of membrane proteins. (B) Study of membrane proteins under different conditions by changing buffer conditions inside/outside the closed vesicle system. (C) Study of mechanical sensor proteins (exampled by PIEZO channel) by preparing vesicles with different sizes.

In combination with affinity grids based on graphene^31^ or graphene oxide^32,33^, our method may provide a novel approach to the structural determination of membrane proteins in their native environment without purification (Figure 3A). Using this approach, vesicles containing target protein can be enriched on grids for cryo-EM imaging and data collection. The purification process will be dramatically simplified, with improved efficiency, enabling the capture of weak binding partners and the study of the interaction network of the target membrane proteins.

## Materials and Methods

### Generation and purification of vesicles

Full-length AcrB was PCR amplificated from the *E*.*coli* genome and subsequently cloned into the pET28a vector with N-terminal His tag. The plasmid was transformed into *E*.*coli* BL21 (DE3) strain and cultured in LB cell culture at 37 °C until OD_600_ (optical density) reached 0.8. The overexpression of AcrB was induced by 0.5 mM isopropyl-β-D-thiogalactoside (IPTG) for 16 h at 20 °C. The cells were harvested at 5,000 g for 20min and resuspended in 40 mL of lysis buffers containing 25 mM Tris-HCl, 150 mM NaCl, and 2 mM MgCl at pH 7.5. The cells were disrupted by the French press under 800pa for 3 cycles. The cell pellet was removed by 6,000 g centrifugation for 30 min and the vesicles were collected under 10,000 g ultra-speed centrifugation for 30 min. The vesicle pellet was washed twice and resuspended into 30 mL buffers. The vesicle samples were homogenized using the French press at 1200 pa for three cycles and filtered through a 0.42 μm Millipore membrane filter. Finally, the vesicle samples were concentrated to approximately 3 mg/mL with the 100 kD Amicon centrifugal filter for further use.

### Density centrifugation of the AcrB vesicles

To further purify the vesicles of uniform size and remove the impurities and irregular vesicles, sucrose density centrifugation was employed. 9mL 30% (W/V) sucrose solution in the lysis buffer was added to a 10mL ultra-speed centrifugation tube and freeze-thawed for 3 cycles between -20°C and 4°C. As previously reported, a smooth sucrose density from 25% to 30% could be generated^34^. The 1mL concentrated vesicle sample was gently added to the top of the top and centrifuged at 50,000g for 14h. 20 fractons were manually aspirated from the centrifugation tube using a pipette, and protein concentration and purity of each fraction were assessed using the UV280 absorption and SDS-PAGE. For negative staining and cyo-EM grids preparation, the fractions were repeatedly diluted with no-sucrose lysis buffer and concentrated using a 100 kD Amicon centrifugal filter to decrease the sucrose concentration to below 0.1%. The final samples were concentrated to around 0.6mg/mL for cryo-EM girds preparation.

### Negative staining

For negative staining grids, 3μL vesicle samples were loaded onto a glow-discharged formvar, carbon-coated copper grid(300mesh, Zhongjingkeyi Technology) and incubated for 45 s. The grids were blotted with filter paper to remove the excess samples and washed with the 2% (w/v) uranyl acetate. Subsequently, the grids were negatively stained with 2% (w/v) uranyl acetate for 45 s and dried with filter paper. The prepared grids were observed and imaged using a Talos L120C electron microscope (Thermo Fisher Scientific) operated at 120 kV with a 4k × 4k Ceta 16M camera. The images were captured with a magnification of 57,000×, corresponding to a pixel size of 2.49 Å on the specimen, and a defocus around -1.5 μm.

### Cryo-EM sample preparation and data collection

Following negative staining results to assess sample quality, uniform purified vesicle samples with great contrast were applied to cryo-grids. We conducted several trials and it was found that the vesicles demonstrated a strong binding preference for carbon film over holes in holey carbon grids. Therefore, the high-concentration sample multiple-loading method was applied. Briefly, 3 μL of high-quality vesicle samples from the previous step were loaded to the glow-discharged Au holey carbon grids(C-flat, 300 mesh, R1.2/1.3) four times, with 45 s incubation at room temperature. Excess samples were quickly blotted with filter paper from the opposite side of the girds to facilitate the vesicles penetrating the holes. The grids were then transformed into a Vitrobot Mark IV (Thermo Fisher Scientifc) and incubated with 3 μL samples again for 45 s at 100% humanity and 4 °C. Then, the grids were blotted with a blot force -1 for 3 s and plunged into liquid ethane by Vitrobot.

The cryo-grids were imaged by a 300 kV Titan Krios G3i electron microscope (Thermo Fisher Scientific) equipped with a K3 Summit direct electron detector (Gatan). Automatic data collection was performed using EPU with 20 eV slit width of the Gatan Imaging Filter (GIF) Bio Quantum in counting mode. Every movie was captured with 4.5 s total exposure time and 40 frames at a calibrated magnification of 81,000× (1.06 Å/pix physical pixel size) to achieve a total dose of 50 e^−^/Å^2^.

### Cryo-EM data processing and model docking

The main steps of data processing were performed in cryoSPARC^25^. A total of 3000 movies were motion-corrected using the patch motion correction with a 7*5 patch. The contrast transfer function(CTF) parameters were calculated by the Patch CTF estimation job.

To accurately pick more particles from the vesicles, multiple strategies were used and combined. The blob picker was first used to locate the particles roughly and the candidates without membrane were removed with the inspect particles picks job due to the low local power value from the intensity. The particles were extracted and 2D classified for several rounds. The featured 2D classes were manually picked as the template input of the template picker. Secondly, the template picker was used to more accurately pick the protein particles from the vesicles. After running the inspect particle picks and 2D classification jobs, clear 2D classes with protein features were chosen manually and their particle locations were input into the Topaz train job^26,27^. Finally, after several rounds of deep learning training, the topaz models were used to precisely pick more AcrB particles from the images. All featured particles from the steps before were merged together, and the repeated particles were removed by the remove duplicate particles job based on their location coordinates.

A total of 56,736 enriched high-quaility particles were re-extracted with a 256 pixel box size after re-centering using the 2D alignment information. After several rounds of 2D classification, the Ab-initio reconstruction was applied to generate an initial map of AcrB in the membrane with 36,454 particles. After a few rounds of homogenous refinement and Heterogeneous refinement were performed, the final 24,025 particles were used to generate the global density map of AcrB with partial vesicle membrane by Non-uniform refinement^35^ at 10 Å. To gain more structural details of the protein, a loose mask covering the AcrB protein region was used to run the Local refinement job with C3 symmetry, and the final density map of AcrB was generated at 5.33 Å.

Every chain of previously reported AcrB cryo-EM structure in native membrane nanoparticles (PDB ID: 6BAJ) was aligned to the density map and merged into a new PDB file in UCSF Chimera^36^. The structural comparison and image drawing were completed using Chimera X^37^.

### Data availability

Cryo-EM density map of AcrB from vesicle has been deposited in the Electron Microscopy Data Bank (accession no. EMD-XXXX). Atomic coordinate has been deposited in the Protein Data Bank (ID codes XXXX). All other data are available from the corresponding authors upon reasonable request.

## Acknowledgments

All cryo-EM data were collected at the Biological Cryo-EM Center at HKUST, generously supported by a donation from the Lo Kwee Seong Foundation. This project is supported by grants from Hong Kong Research Grants Council (ECS26101919, GRF16103321, GRF16102822 and C6001-21EF), Southern Marine Science and Engineering Guangdong Laboratory (Guangzhou) (SMSEGL20SC01), Guangdong Basic and Applied Basic Research Foundation (2021A1515012460), Shenzhen Special Fund for Local Science and Technology Development Guided by Central Government (2021Szvup140), and HKUST start-up and initiation grants.

## Author Contributions

S.D. conceived and supervised the project. H.L. performed all experiments, including vesicle preparation, cryo-EM sample preparation, cryo-EM data collection and processing. H.L and S.D analyzed the data, prepared the figures, and wrote the manuscript.

